# Selective auxin agonists induce specific AUX/IAA protein degradation to modulate plant development

**DOI:** 10.1101/381608

**Authors:** Thomas Vain, Sara Raggi, Noel Ferro, Deepak Kumar Barange, Martin Kieffer, Qian Ma, Siamsa M. Doyle, Mattias Thelander, Barbora Pařízková, Ondřej Novák, Alexandre Ismail, Per Anders Enquist, Adeline Rigal, Małgorzata Łangowska, Sigurd Ramans Harborough, Yi Zhang, Karin Ljung, Judy Callis, Fredrik Almqvist, Stefan Kepinski, Mark Estelle, Laurens Pauwels, Stéphanie Robert

## Abstract

Auxin phytohormones control most aspects of plant development through a complex and interconnected signaling network. In the presence of auxin, AUXIN/INDOLE-3-ACETIC ACID (AUX/IAA) transcriptional repressors are targeted for degradation by the SKP1-CULLIN1-F-BOX (SCF) ubiquitin-protein ligases containing TRANSPORT INHIBITOR RESISTANT 1/AUXIN SIGNALING F-BOX (TIR1/AFB). CULLIN1-neddylation is required for SCF^TIR1/AFB^ functionality as exemplified by mutants deficient in the NEDD8-activating enzyme subunit AUXIN-RESISTANT 1 (AXR1). Here, we report a chemical biology screen that identifies small molecules requiring AXR1 to modulate plant development. We selected four molecules of interest, RubNeddin1 to 4 (RN1 to 4), among which RN3 and RN4 trigger selective auxin responses at transcriptional, biochemical and morphological levels. This selective activity is explained by their ability to promote the interaction between TIR1 and a specific subset of AUX/IAA proteins, stimulating the degradation of particular AUX/IAA combinations. Finally, via a genetic screen using RN4, we revealed that the chromatin remodeling ATPase BRAHMA is implicated in auxin-mediated apical hook development. These results demonstrate the power of selective auxin agonists to dissect auxin perception for plant developmental functions.

## Introduction

The survival and reproductive success of all living organisms depend on their ability to perceive and integrate environmental and internal signals. As sessile organisms, plants have developed strategies to adapt to their surroundings, including an extensive developmental plasticity (1). Plant morphological changes are executed through regulation of hormone levels and signaling (2). The phytohormone auxin is involved in almost all aspects of plant development and adaptation. Auxin perception within the nucleus is mediated by the TRANSPORT INHIBITOR RESISTANT 1/AUXIN SIGNALING F-BOX [TIR1/AFB]-AUXIN/INDOLE-3-ACETIC ACID [AUX/IAA] (TIR1/AFB-AUX/IAA) co-receptor complex (3). The TIR1/AFB1-5 F-box proteins are subunits of the S-PHASE KINASE ASSOCIATED PROTEIN 1-CULLIN 1-F-BOX (SCF)-type E3 ligase and act as auxin receptors (4). Formation of the SCF^TIR1/AFB^-AUX/IAA-auxin complex leads to the ubiquitination of the AUX/IAA transcriptional repressors, targeting them for rapid degradation by the 26S proteasome (4). Removal of AUX/IAAs liberates the auxin response-activating AUXIN RESPONSE FACTOR (ARF) transcription factors from repression (4) and leads to the occurrence of auxin-transcriptional response. There is significant variation in auxin-induced degradation rates among different AUX/IAA proteins, and at least some of this variation is attributable to the specificity in the interactions between the 29 AUX/IAAs and 6 TIR1/AFB F-box proteins in *Arabidopsis* (4–6). Amino acids within and outside the degron domain II (DII) of the AUX/IAA proteins determine the interaction strength of the co-receptor and specify AUX/IAA stability (5–7). The multiplicity of the potential co-receptor assembly is the first element mediating the complexity of the auxin response.

The ubiquitin-proteasome pathway plays an essential role in plant hormone signaling (8–10). Modification of the relevant components by the ubiquitin-like protein, RELATED TO UBIQUITIN/NEURAL PRECURSOR CELL EXPRESSED DEVELOPMENTALLY DOWNREGULATED PROTEIN 8 (RUB/NEDD8), which is catalyzed by a cascade of enzymatic reactions analogous to ubiquitination, is critical for the full activity of the proteasome complex (11). In plants, the CULLINs (CUL1, CUL3, and CUL4) are NEDD8-modified proteins that form multimeric E3 ubiquitin ligase complexes (12). CUL1 acts as a scaffold within the SCF-type E3 ligases and neddylation states of CUL1 are essential for the ubiquitin ligase activity of the SCF complex (13). Loss of components of the neddylation pathway, such as the NEDD8-activating enzyme subunit AUXIN RESISTANT 1 (AXR1), reduces the response to several phytohormones including auxin (14–17).

To understand how auxin perception mediates multiple aspects of plant development, we established an AXR1-dependent developmental defect-based chemical biology screen. Using this approach, we identified new small synthetic molecules that selectively promote SCF^TIR1/AFB^-AUX/IAA co-receptor assembly, allowing local and precise modulation of auxin signaling pathways. Furthermore, these synthetic selective agonists possess the ability to identify and distinguish the molecular players involved in different aspects of auxin-regulated development, thereby dissecting the diversity of auxin action. We demonstrated this by employing these agonists to reveal different roles for specific AUX/IAA proteins during lateral root and apical hook development. In particular, the use of the selective auxin agonist RN4 revealed a new role for the chromatin remodeling ATPase BRAHMA in apical hook development.

## Results

### The rubylation/neddylation pathway is required for RubNeddins (RNs) to alter seedling development

In order to address the complexity of auxin response, we established a chemical biology screen to isolate synthetic molecules targeting the NEDD8-mediated signaling pathway in *Arabidopsis* (Fig. S1*A* and *B*). We reasoned that some of these molecules might also target the auxin signaling pathway (Fig. S1*A*). This strategy is complementary to previous ones aiming at isolating auxin-related small molecules (18–19). Compounds affecting auxin-related developmental processes such as primary root growth, hypocotyl elongation and gravi- or photo-tropism responses in wild type but not in *axr1-30* seedlings were selected (Fig. S1*B*). This screening strategy, based on differential effects upon the two genetic backgrounds (Col-0 wild type versus *axr1-30*), was essential to filter out chemical activities with general impacts on seedling growth. We hypothesized that a small molecule for which activity was dependent on the AXR1 signaling machinery could be recognized by one or several TIR1/AFB-AUX/IAA coreceptor complexes. Out of 8,000 diverse compounds (Chembridge), we identified 34 small molecules (4.25 ‰) that selectively affected the growth of wild type compared to *axr1-30* seedlings, indicating that their activities require a functional AXR1. Four molecules, named RubNeddin (RN) 1 to 4, were ultimately selected as they showed a dose-dependent activity and a high potency on wild type seedling development in the micromolar range (Fig. S1*C-E*). In detail, RN1 activity decreased lateral root number and primary root length, but increased hypocotyl length and adventitious root formation (Fig. 1*A* and *B*, Fig. S2*A*). RN2 application resulted in the inhibition of primary root growth and lateral root formation, without affecting hypocotyl length (Fig. 1*A* and *C*). RN3 promoted the number of lateral roots (Fig. 1*A* and *D*). RN4 activity increased hypocotyl elongation and inhibited lateral root formation (Fig. 1*A* and *E*). Overall, these structurally similar compounds triggered specific morphological changes in wild type, while *axr1-30* was resistant to these effects, demonstrating that they require a functional RUB/NEDD8 signaling pathway.

**Figure 1.**
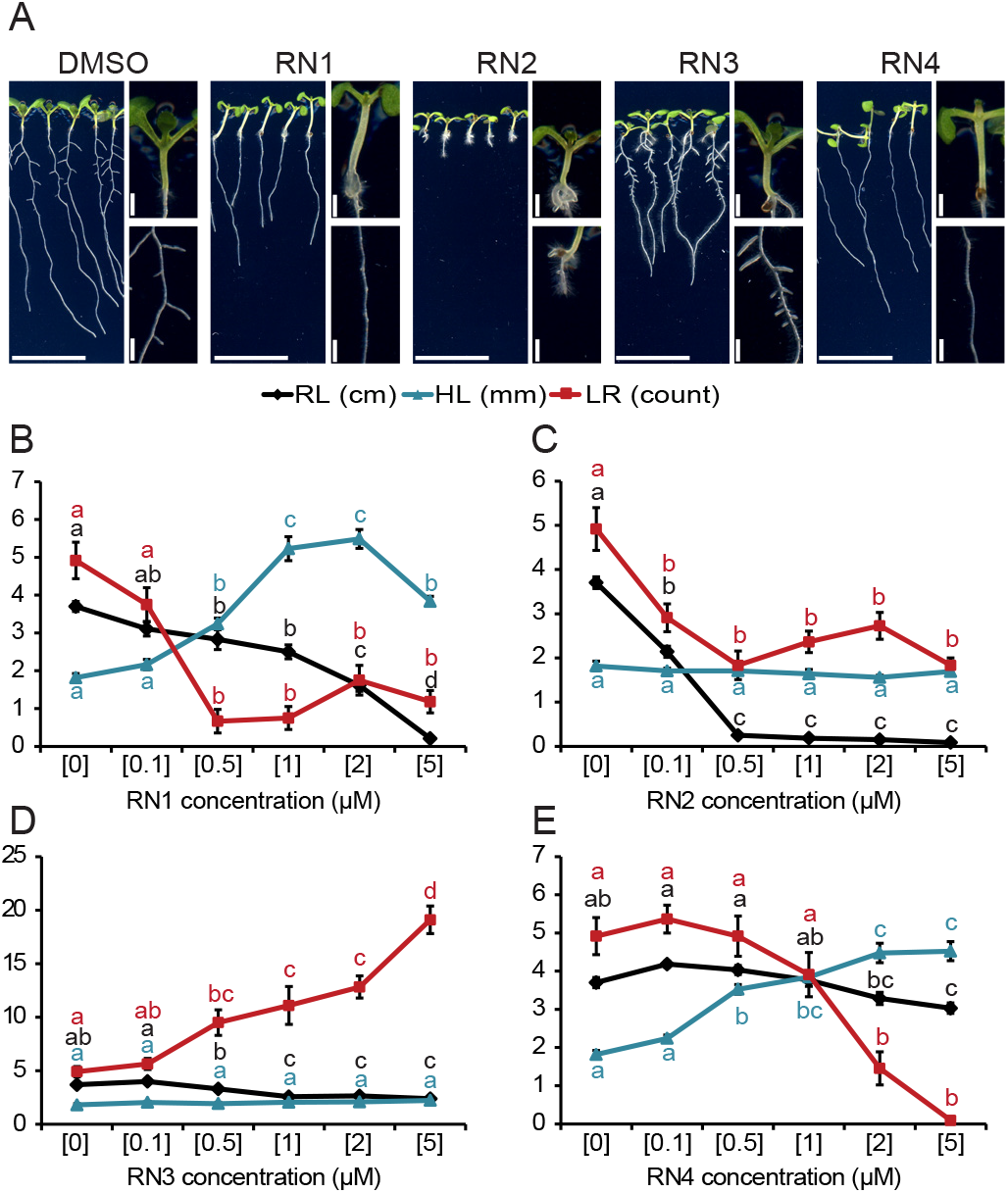
Four RN chemicals trigger different morphological changes. (A) Col-0 seedlings were grown on RN-supplemented media for eight days. DMSO was used as control. Images display the effects of the RN at a representative concentration: RN1: 2 μM; RN2: 0.5 μM; RN3: 2 μM; RN4: 5 μM. (B-E) RN1 (B), RN2 (C), RN3 (D) and RN4 (E) selectively affected primary root length (RL), hypocotyl length (HL) and the number of lateral roots (LR). Statistics were performed using ANOVA and Tukey’s test. Means ± SEM are shown, n = 10 seedlings for each concentration of the dose response, different letters are displayed for p-value < 0.05. Scale bars indicate 1 cm (A). Concentrations in μM are indicated in brackets (B-E).

### The RNs act as developmental regulators in several land plants

We then analyzed RN effects on *Populus* (poplar) and *Physcomitrella patens* (moss) to test their activities on several land plants. RN1, which induced hypocotyl elongation and promoted adventitious root formation, and RN3, which increased lateral root number in *Arabidopsis*, were applied to three different lines of poplar explants (Fig. S2*B-D*). The poplar lines were selected for their different rooting abilities; T89 is an easy rooting hybrid while SwAsp 19 and 35 have a low rooting capacity even when treated with indole-3-butyric acid (IBA), an auxin commonly used as a rooting agent. Interestingly, both RN1 and RN3 promoted adventitious root formation preferentially in the SwAsp lines. Next, the effects of the RNs were investigated in moss and compared to those of IAA (Fig. S3). Similar to IAA, most of the RNs inhibited caulonemal colony outgrowth (Fig. S3*A*). The RN-induced effects on shoots were more diverse. At the tested concentrations, while no effect of RN1 was observed, application of RN2 caused a clear increase in shoot length, RN3 treatment resulted in thinner leaves and RN4 slightly reduced shoot size (Fig. S3*B*). At low concentration, IAA increased the number of buds/shoots per colony after one week (Fig. S3*C*), while it reduced bud/shoot formation after two weeks regardless of the concentrations tested (Fig. S3*D*).

This dual effect of IAA was mimicked by RN4. RN1 and RN3 treatment resulted mainly in an increase of the bud/shoot number per colony after one week and RN2 only reduced bud/shoot formation after two weeks. These results demonstrate that the activities of the RNs are mediated by pathways present in several species.

### The RNs function as prohormones

RN1, RN3 and RN4 share structural similarities with previously described prohormones (19–20). Since prohormones are hydrolyzed *in vivo* to release the active hormone moieties (21), we examined the potential metabolism of the RN compounds in liquid treatment media and *in planta* (Fig. S4). In RN-supplemented MS media without plants, negligible concentrations of free acids were detected at the 0 h time point, except for 2,4-dichlorophenoxyacetic acid (2,4-D) originating from RN2 and 2,4,5-trichloroacetic acid (2,4,5-T) from RN3 (Fig. S4*D*). Importantly, in these plant-free media, no obvious degradation of RN compounds was observed 24 h after treatment. However, in the presence of seedlings, higher levels of the corresponding free acids, 2,4-D, 2,4,5-T and RN4-1, were found after 24 h in the media treated with RN1, RN3 and RN4, respectively, although the level of 2,4-D in RN2-treated media was not changed (Fig. S4*D*). As expected, in *Arabidopsis* seedlings treated by the RNs for 24 h, all free acids were detected in the range from 0.4 to 2% relative to the levels of the corresponding RNs (Fig. S4*E*). These results imply that even though the RN compounds are fairly stable in liquid media, their biological activities might result from their metabolism *in planta* to the free acids 2,4-D (RN1 and RN2), 2,4,5-T (RN3) and RN4-1 (RN4), all of which are known to possess auxinic activity. To address this possibility, we performed a structure activity relationship (SAR) analysis by comparing the effects of various RN analogues, 2,4-D, 2,4,5-T and RN4-1 on plant development and we evaluated their ability to modulate the expression pattern of the auxin responsive promoter *DR5* in seedlings of p*DR5*::GUS (22) (Fig. S5).

The SAR analysis indicated that the absence of chlorine at position C2 in the 2,4-D substructure of RN1 (analog RN1-1) or the complete loss of the 2,4-D moiety (analog RN1-2) significantly reduced the effects of RN1 on plant development (Fig. S5*A* and *E*), implying that the 2,4-D substructure is important for RN1 activity. However, 2,4-D itself induced phenotypes distinct from RN1, suggesting the activity of this RN requires both 2,4-D and non-2,4-D substructures. Neither RN1 nor the RN1 analogues visibly altered the p*DR5*::GUS expression pattern compared to the DMSO control. Modification of the 2,4-D core structure in RN2 (analog RN2-2) abolished its potency, whereas analogs displaying a side chain modification (RN2-1 or RN2-3) were as potent as RN2 (Fig. S5*B* and *F*), indicating that the activity of RN2 is most probably attributable to the release of 2,4-D from the compound after dissolution in the growing media. Like RN2, none of the RN2 analogues visibly altered the p*DR5*::GUS expression pattern compared to the DMSO control. RN3 mainly promoted lateral root number, while its effect on primary root elongation was mild (Fig. 1*D*). Analogs RN3-2 and RN3-3, with modifications on the phenylpiperazine side chain, behaved similarly to RN3 (Fig. S5*C*, *G* and *H*). However, removal of the whole side chain from RN3, generating 2,4,5-T, abolished its positive effect on lateral root number and introduced a strong inhibitory effect on primary root length (Fig. S5*H*), suggesting a difference in potency between the two compounds. Moreover, the activity of RN3 was significantly compromised by disruption of the substructure of 2,4,5-T (analog RN3-1) via loss of the three chlorines (Fig. S5*C*, *G* and *H*). These results suggest that the 2,4,5-T substructure is critical for RN3’s potency. Further comparisons using analogs only differing in the number of chlorines on the 2,4,5-T substructure, such as between RN3-2, RN3-4 and RN3-6, or between RN3-3, RN3-5 and RN3-7, indicated that C5 chlorination of the 2,4,5-T moiety is crucial for RN3’s selective activity. Intriguingly, while RN3 did not alter the p*DR5*::GUS expression pattern compared to the DMSO control, fluorination of the phenyl in RN3 induced p*DR5*::GUS expression in some cases (analog RN3-3 compared to RN3-2), while reducing it in other cases (analogs RN3-5 and RN3-7 compared to RN3-4 and RN3-6, respectively) (Fig. S5*C*). These results reinforce the importance of C5 chlorination of the 2,4,5-T moiety for the selective activity of RN3.

We showed that RN4 releases the free acid RN4-1 *in planta* (Fig. S4*D* and *E*), possibly by hydrolysis. RN4-1 contains a bromo group, an electron-withdrawing substituent, which can give rise to a high auxinic activity (23) and as expected, this compound strongly induced p*DR5*::GUS expression (Fig. S5*D*). In contrast, RN4 treatment did not promote p*DR5*::GUS expression in the root (Fig. S5*D*). However, while RN4-1 significantly enhanced hypocotyl elongation, it was not as potent in this regard as RN4 (Fig. S5*D* and *I*). Comparison of the effects of modifications of the RN4-1 substructure (analog RN4-2) and of the hydroxymethylphenylamine substructure (analog RN4-10) of RN4 indicate that while the intact auxinic RN4-1 moiety is indispensable for RN4’s effect on the hypocotyl, the non-auxinic side chain is also required to induce maximal hypocotyl elongation (Fig. S5*D* and *I*). Further comparison between RN4-2 and RN4 as well as their free acids (RN4-3 and RN4-1, respectively) highlight the key contribution of the bromophenoxy methylation to the selective activity of RN4 on hypocotyl rather than primary root (Fig. S5*DI* and *J*). Consistent with the SAR results, even though RN4-2 shows a bipartite structure, it was still able to induce p*DR5*::GUS expression (Fig. S5*D*). RN4-10, in which the non-auxinic moiety of RN4 is modified, induced p*DR5*::GUS expression slightly more than RN4 (Fig. S5*D*). We also designed RN4 analogs with predicted low hydrolysis capacity (RN4-4, RN4-8, RN4-9, and RN4-11). As expected, none of these analogs could induce hypocotyl growth (Fig. S5*D* and *I*), indicating that the typical bipartite prohormone structure of RN4 is important for its effect on hypocotyl elongation and that hydrolysis is required to liberate this activity. Moreover, except for RN4-9, these compounds could not induce p*DR5*::GUS. Interestingly, the analog RN4-11, generated by methylation of RN4 on the amide bond, inhibited primary root elongation without affecting hypocotyl length (Fig. S5*J*). As the predicted corresponding free acid RN4-1 did not reduce primary root length, this result indicates that the full, non-hydrolyzed RN4 structure possesses additional auxin-like activity.

Overall, we showed that RN1, RN3 and RN4 function as prohormones, being metabolized *in planta* to release more potent auxin agonists, while the effects of RN2 are most likely due to its degradation to 2,4-D. However, our SAR results also suggest that the non-hydrolyzed forms of RN1, RN3 and RN4 display additional auxin-like effects and therefore might themselves act as selective auxin agonists.

### The RNs act as selective auxin agonists

AXR1 is a component of the neddylation pathway targeting, among others, the CUL proteins (11). To determine which CUL proteins might be involved in mediating the effects of each RN, we tested their potency on the loss of function *cul1-6*, *cul3a*/*b*, and *cul4-1* mutants. We limited these tests to RN1, RN3, and RN4 as we showed that RN2 activity is most probably due to its *in vitro* cleavage into 2,4-D, an already well described synthetic auxin. All three tested RNs had a lesser effect on the *cul1-6* mutant than on other CUL mutant lines (Fig. 2*A*). This result indicates that the three RNs function at the level of or upstream of CUL1. Given that signaling pathways mediated by AXR1 and CUL1 converge at the SCF complex and that the chemical structures and activities of the three RNs are related to auxin, we hypothesized that auxin receptor F-box proteins might also be required for RN activities. To test this, we examined *tir1* single and *tir1/afb* multiple mutants and found that the RN-induced phenotypes were strongly reduced when the compounds were applied on *tir1-1* and *tir1-1afb1*-3*afb3-4* (24–25) (Fig. 2*B*). Thus, a functional SCF^TIR/AFB^ complex is essential for the effects of the RNs. To further confirm this result, we tested the effect of co-treatment of the compound auxinole (26), an auxin antagonist specific for SCF^TIR1/AFB^, together with each of the three RNs or the endogenous auxin IAA in the wild type. The RN-induced phenotypes were inhibited by auxinole (Fig. 2*C*), demonstrating that auxin co-receptor complex formation is essential for RN activities.

**Figure 2.**
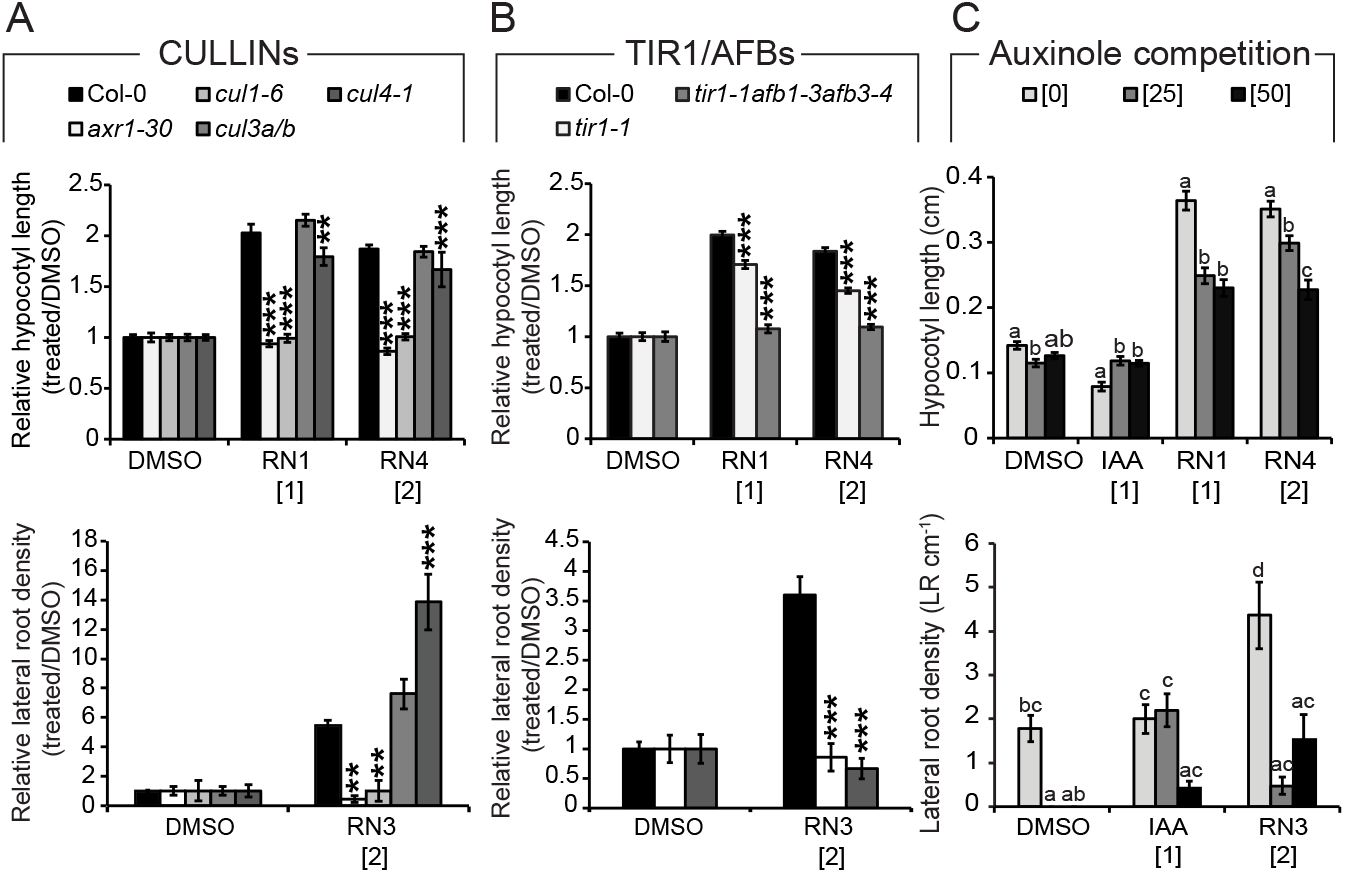
RN-induced phenotypes require the formation of a functional auxin-SCF^TIR1/AFB^ complex. (A-C) Hypocotyl length (upper charts) and lateral root density (lower charts) were measured for wild type (Col-0) and mutant seedlings grown on media supplemented with RN compounds for seven days. DMSO was used as control. (A) Mutants *axr1-30* and *cul1-6* were less sensitive to the representative RN-induced phenotypes than Col-0, while *cul3a/b* and *cul4-1* were still affected. (B) Mutants *tir1-1* and *tir1-1afb1-3afb3-4* were less sensitive to the RN-induced phenotypes than Col-0. (C) The RN-induced phenotypes were inhibited by auxinole application in Col-0. Statistics were performed using ANOVA and Tukey’s test. Means ± SEM are shown, n = 30 seedlings across 3 independent replicates, p-value: \*\**P* < 0.01, \*\*\**P* < 0.001 (A-B) or different letters are displayed for p-value < 0.05 (C). Concentrations in μM are indicated in brackets.

Next, we employed a molecular modeling strategy to explore the possible interactions of the RNs with the DII degron of AUX/IAA7 in the auxin-binding pocket of TIR1. Docking experiments validated that the physical property of the auxin-binding pocket was promiscuous enough to accommodate the potential steric hindrance of RN1, RN3, or RN4 (Fig. 3*A-C*; Movie S1). The calculated free energies (ΔG) of binding also revealed thermodynamic stability for the three RNs inside the auxin pocket of TIR1 (Fig. 3*A-C* and Fig. S6*A*). The positive control IAA was able to bind TIR1 with a Δ*G*_(IAA-TIR1)_ of −11.68, whereas the negative control Tryptophan (Trp) was not, with a Δ*G*(Trp-TIR1) of 63.34 (Fig. S6*A*). Among the RN analogs, RN4-1 and RN4-2 showed stronger thermodynamic stability compared to IAA. RN2 and the inactive analog RN4-8 could not dock inside the auxin-binding site to stabilize TIR1 (Fig. S6*A*). This last result confirmed once again that RN2 activity is most likely due to its cleavage into 2,4-D.

**Figure 3.**
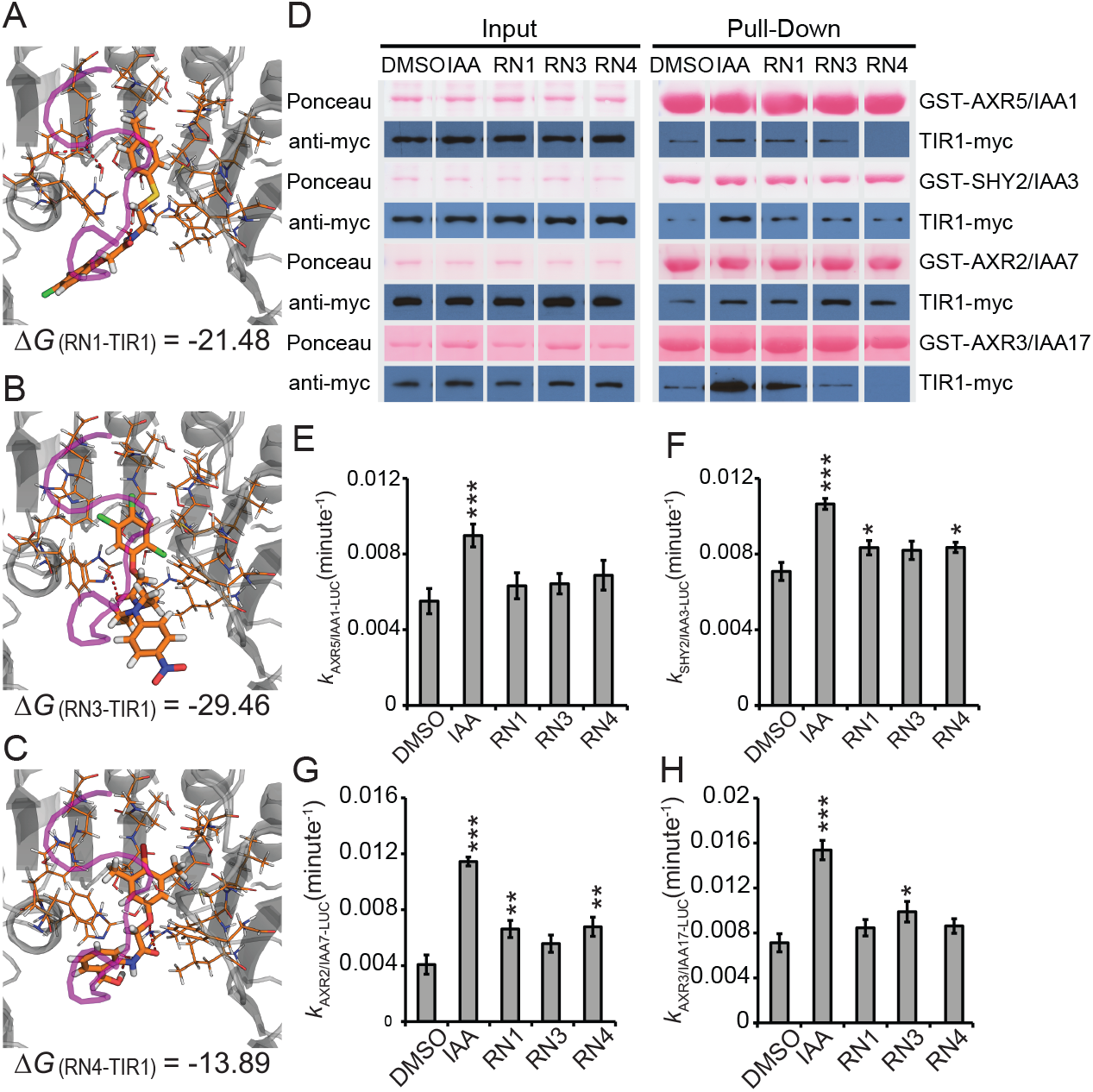
RN3 and RN4 act as selective agonists of auxin. (A-C) The RNs showed different thermodynamic stabilities from the calculated free energies (ΔG). RN1 (A), RN3 (B) and RN4 (C) were sterically favorable for the binding of the AUX/IAA7 DII degron. TIR1 is presented in gray and the AUX/IAA7 DII degron, which was included afterwards to observe any conflict with the RNs, is in purple. Thermodynamic stability was computed within the TIR1 auxin binding pocket and the most stable conformation(s) is represented. (D) The potential of the RNs (at 50 μM) to promote the formation of the co-receptor complex was performed using *in vitro* translated TIR1-myc and recombinant GST-AUX/IAAs. Depending on the GST-AUX/IAA translational fusion used for the *in vitro* GST pull-down, the RNs selectively increased the recovery of TIR1-myc. (E-H) AUX/IAA degradation was assayed *in planta* using *Arabidopsis* lines constitutively expressing different AUX/IAA-LUCs in the presence of RNs at 50 μM. Effects of the RNs on the *in vivo* degradation rate *k* of AXR5/IAA1-LUC (E), SHY2/IAA3-LUC (F), AXR2/IAA7-LUC (G) or AXR3/IAA17-LUC (H) translational fusions. Statistical analyses were performed using the Student’s t-test. Means ± SEM are shown, n = 30 seedlings across 5 independent replicates, p-value: *P* < 0.1, \**P* < 0.05, \*\**P* < 0.01, \*\*\**P* < 0.001.

To experimentally confirm the binding of the RNs within the auxin co-receptor complex, we tested their ability to promote the interactions between TIR1 and AUX/IAA proteins using *in vitro* pull-down assays. First, TIR1-myc protein purified from wheat germ extract and four different GST-AUX/IAA proteins were used (27–29). IAA stimulated the interaction of TIR1-myc with all AUX/IAAs tested (Fig. 3*D*). All three RNs stimulated the recovery of TIR1-myc in complex with GST-SHY2/IAA3 or GST-AXR2/IAA7 to a similar extent (Fig. 3*D*). In the case of GST-AXR5/IAA1, RN1 stimulated the interaction with TIR1-myc while RN3 had little effect and surprisingly, RN4 decreased the basal interaction (Fig. 3*D*). When GST-AXR3/IAA17 was used as bait, RN1 strongly promoted the interaction with TIR1-myc, while RN3 had no effect and again, RN4 reduced the basal interaction. These data imply that RN3 and RN4 are able to selectively promote the interactions between specific TIR1 and AUX/IAA protein combinations in this system, while RN1 and IAA promoted each interaction, as shown previously for IAA (27–29). To test that these effects on TIR-AUX/IAA complex formation were not dependent on metabolism of the RN compounds in the wheat germ extract, we next performed a complementary pull-down experiment using insect cell-expressed TIR1 (as a His-MBP-FLAG-TIR1 fusion protein) with bacterially-expressed GST-AXR2/IAA7 or GST-AXR3/IAA17 in the presence of the RNs or the RN4 degradation product RN4-1 (Fig. S6*B*). In this system, the RNs again promoted selective interactions between TIR1 and AXR2/IAA7 or AXR3/IAA17, this time in the absence of potential plant hydrolases (in insect cells). Importantly, the promotion and inhibition of TIR1 interaction with AXR2/IAA7 and AXR3/IAA17 respectively by RN3 and RN4 were identical in the two *in vitro* systems. Moreover, the degradation product RN4-1 behaved differently from RN4, by not promoting the interaction between TIR1 and AXR2/IAA7 and slightly promoting the interaction between TIR1 and AXR3/IAA17. The fact that the degradation product RN4-1 behaves differently from RN4 *in vitro* might explain these compounds’ different activities *in vivo*. These data demonstrate that RN3 and RN4 are able to selectively promote the interactions between TIR1 and certain AUX/IAA proteins, while IAA strongly promotes each interaction, as shown previously (27–29). Hence, our results suggest that RN3 and RN4 are not just prohormones, but also act consistently as selective auxin agonists in two different *in vitro* experimental conditions and their effects on plant development may therefore be attributable to selective auxin agonistic activity.

To test whether the RNs might also act as selective auxin agonists *in planta*, we assayed their potency in promoting the *in vivo* degradation of the AUX/IAA proteins. In a one-hour time course, IAA was able to significantly increase the degradation rate of the four tested AUX/IAA-LUCIFERASE (LUC) proteins, while the RNs had different potency depending on the AUX/IAA proteins used (Fig. 3*E-H* and Fig. S6*C-F*). Therefore, the RN molecules are able to act as selective auxin agonists both *in vitro* and *in vivo*, but it should be noted that the specificity of the interactions seems to be dependent on the experimental conditions, as the predicted behavior of AUX/IAA proteins based on their sensitivity to RN3 and RN4 in our *in planta* LUC assays did not always match that in our *in vitro* pull-down assays. While the conditions tested *in vivo* reflect RN capacity to enhance the interactions of the different SCF^TIR1/AFB^-AUX/IAA co-receptors within a complex molecular surrounding, those tested *in vitro* reflect the interactions in much simpler conditions. Nonetheless, our results imply that altering interaction affinity within each co-receptor complex with selective auxin agonists might modulate a multitude of specific plant development aspects.

### RN3 and RN4 induce selective early transcriptional responses

The *in vitro* assays indicated that RN3 and RN4 are the most selective auxin agonists, showing different effects on different AUX/IAA proteins. Moreover, RN3 and RN4 induced distinct developmental processes, particularly on lateral root development. While RN3 enhanced the density of lateral roots by increasing their number without affecting primary root length in the wild type, RN4 inhibited lateral root development (Fig. 1). As these RNs promoted fast degradation of AUX/IAA proteins fused to LUC, we decided to investigate how their activities fine-tuned events downstream of coreceptor complex formation. To this end, we performed transcriptome-wide expression profiling of *Arabidopsis* cell suspension cultures treated with IAA, RN3 and RN4, to characterize the early transcriptional responses induced by these compounds (*Dataset 1*). Analysis of the differentially expressed genes (DEGs) revealed subsets of genes that were up- or down-regulated specifically by one, two or all three chemical treatments (Fig. S7*A* and Table S1). Among the early auxin-responsive genes identified, *AXR5/IAA1, IAA2, SHORT HYPOCOTYL 2 (SHY2)/IAA3* and *IAA30* were significantly up-regulated by IAA, RN3 and RN4 (Fig. 4*A* and Table S1). *IAA5* and *IAA16* expressions were induced specifically by IAA and RN3, while *IAA10* and *IAA29* expressions were upregulated selectively by IAA and RN4, revealing some differences between RN3 and RN4 in their capacity to induce early-responsive *AUX/IAA* genes. In total, 121 genes were differentially up-regulated by IAA, RN3 and RN4, such as *LATERAL ORGAN BOUNDARIES-DOMAIN 16 (LBD16), BASIC HELIX-LOOP-HELIX 32 (BHLH32), PINOID-BINDING PROTEIN 1* (*PBP1*) and *PIN-FORMED 3* (*PIN3*) (30–33) (Fig. 4*A*), confirming the potential of the RNs to modulate auxin-related developmental processes. The genes *CINNAMATE 4 HYDROXYGENASE* (*C4H*), *TRANSPARENT TESTA 4* (*TT4*), *TT5, DEHYDRATION RESPONSE ELEMENT-BINDING PROTEIN 26* (*DREB26*) and *EARLY-RESPONSIVE TO DEHYDRATION 9* (*ERD9*) were commonly up-regulated by IAA and RN3 but not by RN4. These five genes are known to be tightly regulated in a tissue-specific and auxin-dependent manner to modulate lateral root density and architecture (34–38). Among the genes commonly regulated by IAA and RN4 but not RN3, we identified *MYELOBLASTOSIS 77* (*MYB77*) and *BREVIX RADIX* (*BRX*) transcription factors, which have been shown to control lateral root formation in an auxin-dependent manner (39–40). These results correlate with the differential effects of RN3 and RN4 on lateral root development. Taken together, these data demonstrate the potential of RN3 and RN4 to specifically identify auxin-responsive genes involved in defined developmental processes such as lateral root formation. Overall, we showed that RN molecules are able to selectively trigger specific auxin perception machinery, inducing expression of specific sets of gene, and resulting in distinct developmental traits.

**Figure 4.**
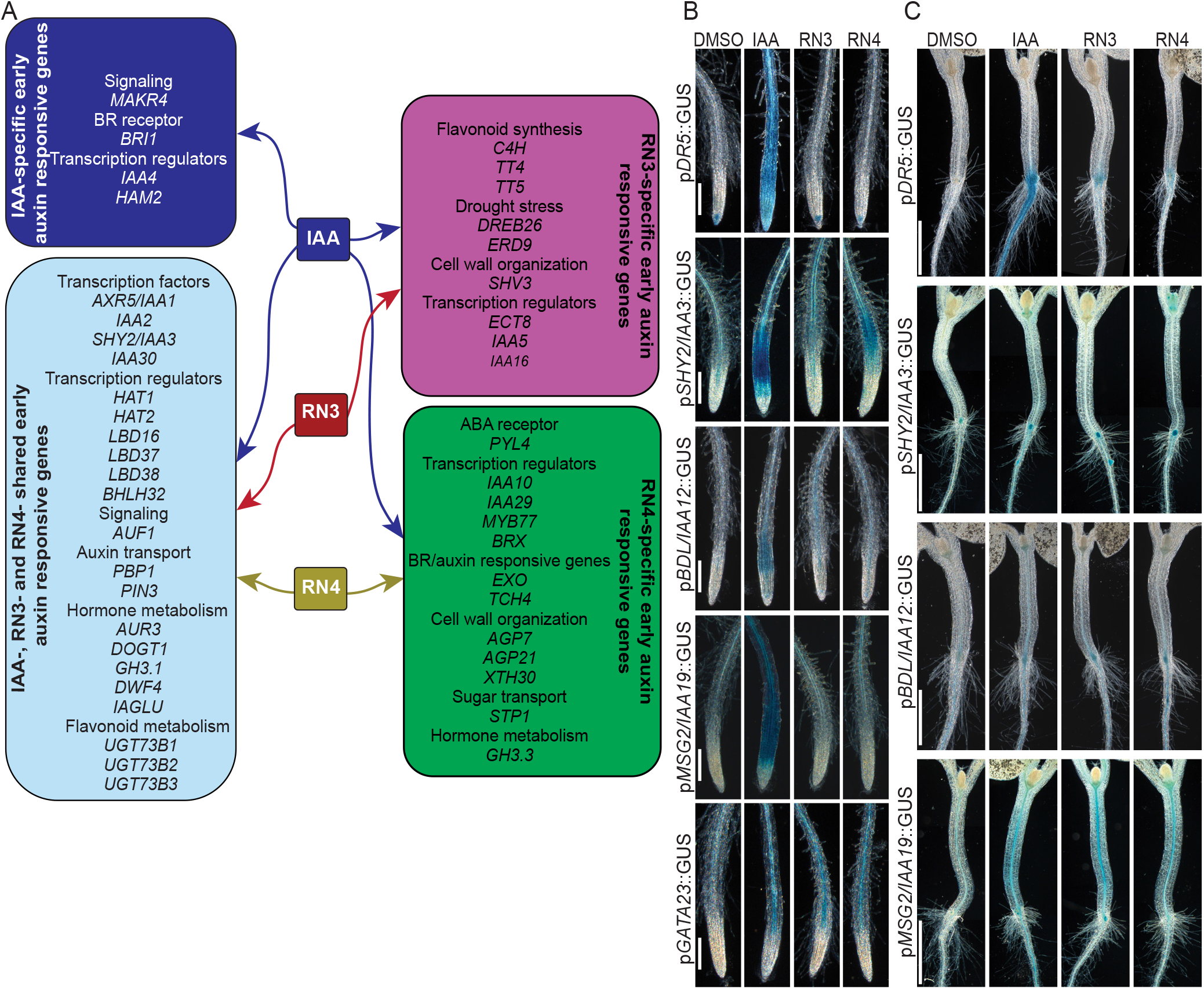
RN3 and RN4 activate independent auxin responses. (A) Selected sets of upregulated genes in cell culture representing: IAA-specific-induced genes (dark blue); IAA-, RN3- and RN4-induced genes (light blue); IAA- and RN3-specific-induced genes (lilac) and IAA- and RN4-specific-induced genes (green) (see *Dataset 1* for the complete list of genes and Table S1 for fold induction values of the selected genes). (B-C) Five-d-old seedlings expressing p*DR5*::GUS, *pSHY2/IAA3*::GUS, pBDL/IAA12:: GUS, *pMSG2/IAA19*::GUS or *pGATA23:*:GUS transcriptional fusions treated with IAA, RN3 and RN4 at 10 μM for 16 h. DMSO was used as control. The RN-induced expression pattern was promoter-dependent and restricted to tissues where the auxin response was observed. (B) Representative primary roots after GUS staining. (C) Representative hypocotyl-root junctions after GUS staining. Scale bars indicate 100 μm (B) and 1 mm (C).

### RN3 and RN4 induce specific subsets of auxin responsive promoters

We further investigated the abilities of RN3 and RN4 to selectively induce later auxin responses using various auxin-responsive reporter lines after 45 min, 5 h or 16 h of RN treatment. We found that neither the auxin-responsive reporter p*DR5*::GUS nor the indicator of nuclear auxin perception p*35S*::DII-Venus (41) showed any response to RN treatment in the primary root (Fig. 4*B*, Fig. S7*B* and *D*). However, in the root-hypocotyl junction, the expression of p*DR5*::GUS was promoted by either longer treatment (24 h) or higher concentration (50μM) of RN3 or RN4 (Fig. 4*C* and Fig. S7*C*). For other auxin-responsive reporter lines tested, RN3 and RN4 induced auxin-responsive promoter expression patterns that partially overlapped with those induced by IAA (Fig. 4*B, C*). In the primary root, the RN compounds induced *pSHY2/IAA3*::GUS and *pBODENLOS(BDL)/IAA12*::GUS expression with different patterns compared to that induced by IAA, but did not stimulate *pMASSUGU2(MSG2)/IAA19*::GUS expression (Fig 4*B*). Both compounds also promoted the expression of p*GATA23:*:GUS, a marker of lateral root founder cell identity (42). RN4 additionally induced p*SHY2/IAA3*::GUS expression in the hypocotyl and the shoot apical meristem (SAM) (Fig. 4*C*). In contrast to the primary root, RN3 and RN4 induced p*MSG2/IAA19*::GUS expression in the hypocotyl (Fig. 4*C*), although only RN4 induced hypocotyl elongation (Fig. 1*B*). Our data indicates that RN3 and RN4 are able to induce specific auxin-regulated promoters, which might be responsible for their selective activities on plant development. Indeed, these RNs activate some but not all modules of the auxin signaling pathway within the same tissue, confirming their selective auxin agonist activities.

A summary of the results obtained for the four RNs is presented in Table S2. In particular, RN3 and RN4 behave as auxin agonists which selectively promote or inhibit AUX/IAA degradation in a reproducible manner leading to specific transcriptional regulation and developmental outputs.

### AUX/IAA sensitivity to RN3 and RN4 *in planta*

We hypothesized that as the RN molecules show selectivity towards the auxin coreceptor complex, they might help to dissect specific functions of individual AUX/IAAs in distinct developmental processes. One approach to dissect which particular AUX/IAAs must be degraded to regulate specific developmental processes could be to investigate the responses of AUX/IAA gain-of-function mutants to auxin treatment; however, such a genetic approach could prove problematic due to high redundancy among the AUX/IAAs. As a potentially more effective alternative, we challenged such mutants with the specific auxin analogs RN3 and RN4.

We first focused on lateral root development as RN3 and RN4 had opposite effects on this process (Fig. 1*D* and *E*). Furthermore, based on our transcriptomic analysis, RN3 and RN4 induce different sets of IAA-responsive genes that are known to be involved in the regulation of lateral root development (Fig. 4*A*). We therefore investigated the sensitivities of 8-d-old seedlings of AUX/IAA gain-of-function mutants *axr5-1/iaa1, axr2-1/iaa7, shy2-2/iaa3* and *solitary root (slr-1)/iaa14* to treatments of RN3, RN4 and 2,4,5-T, RN4-1 (as controls) with regards to lateral root development. Most of the AUX/IAA gain-of-function mutants tested showed similar phenotypes as published previously in control conditions (28, 43–45). However, *axr2-1/iaa7* developed similar lateral root density as the wild type, in contrast to the increased density that has been shown previously (46) (Fig. S8*A*).

We tested the sensitivities of these gain-of-function mutants to RN3, which increases lateral root density in the wild type (Fig. *5A* and Fig. S8*A*) and found that the mutants were sensitive to this effect (Fig. 5*A* and Fig. S8*A*), with the exception of *slr-1/iaa14* (Fig. 5*A* and Fig. S8*A*). It is important to note that in *shy2-2/iaa3*, which showed hypersensitivity to RN3, this compound mainly induced the slight emergence of lateral root primordia rather than the emergence of well-developed lateral roots. These data suggest that apart from SLR/IAA14, the AUX/IAAs we tested are not required for the stimulatory activity of RN3 on lateral root density. As a control for RN3, we also tested its degradation product 2,4,5-T at 0.2 μM, because this concentration induces a similar lateral root phenotype as RN3 treatment at 2 μM (Fig. S8*C*). Importantly, mutant sensitivity to 2,4,5-T was different to that observed with RN3 (Fig. 5*A* and Fig. S8*A*), suggesting that these compounds act in different ways to promote LR formation.

**Figure 5.**
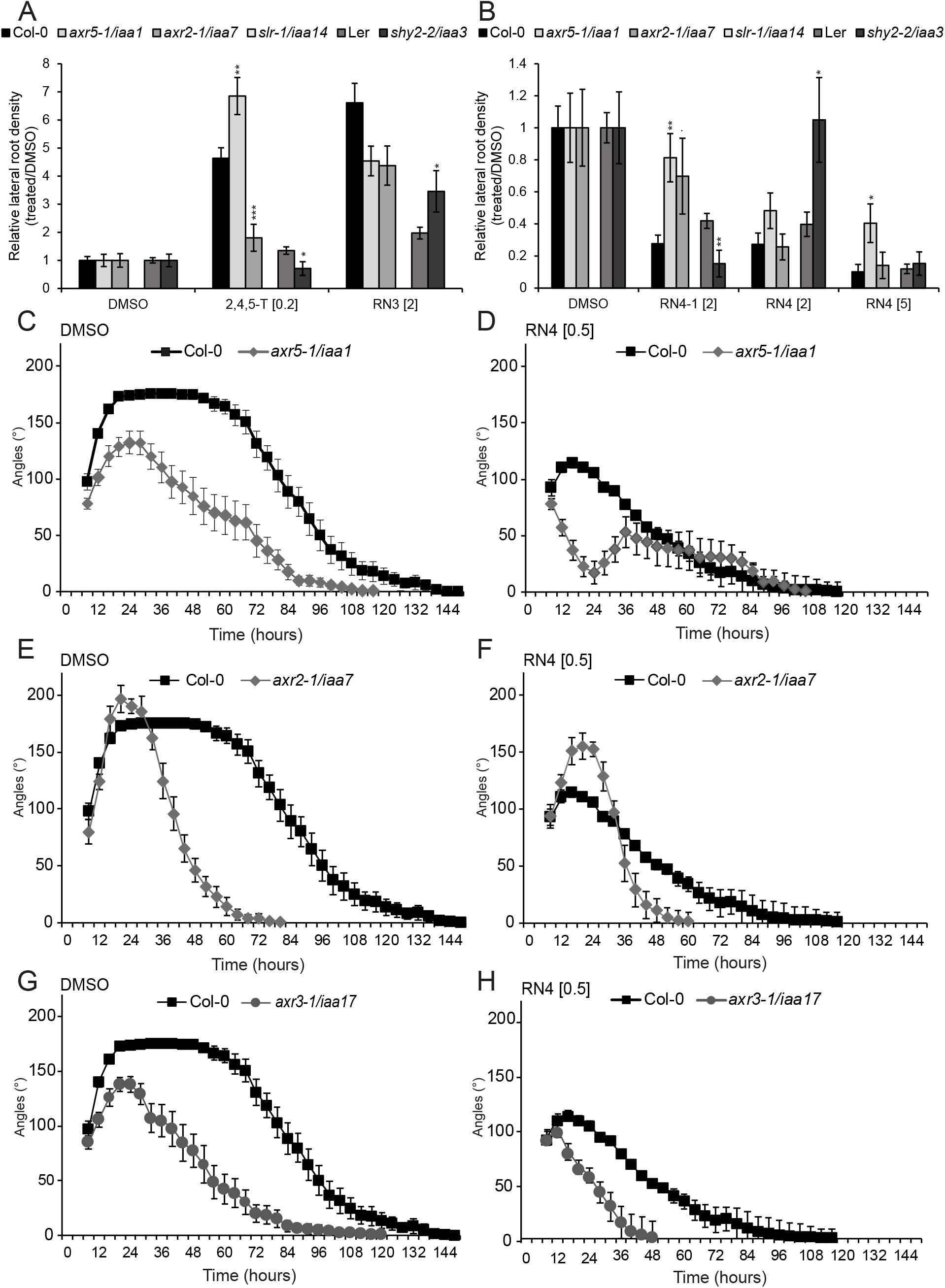
RN-induced phenotypes require the degradation of specific AUX/IAAs. (AB) Relative lateral root density (treated/DMSO) was measured for gain-of-function mutants *axr5-1/iaa1, axr2-1/iaa7, slr-1/iaa14* and *shy2-2/iaa3* and their respective wild type grown on media supplemented with RN3 (A) and RN4 (B) and their respective degradation products for eight days. DMSO was used as control. (C-H) Gain-of-function mutants *axr5-1/iaa1* (C, D), *axr2-1/iaa7* (E, F) and *axr3-1/iaa17* (G, H) were grown in the dark on DMSO (C, E, G) and RN4 (D, F, H) supplemented media for 6 days. Measurement of apical hook angle was performed every three hours. Statistical analyses were performed using ANOVA and Tukey’ s test to compare the effect of RN3 (A) and RN4 (B) and their degradation products on *axr5-1/iaa1, axr2-1/iaa7* and *slr-1/iaa14* relative to the chemical effect on Col-0. Statistical analyses were performed using Student’s t-test to compare the effect of RN3 (A) and RN4 (B) and their degradation products on *shy2-2/iaa3* relative to the chemical effect on Ler. Means ± SEM are shown, n > 20 seedlings across 3 independent replicates, p-value: *P*< 0.1, \**P* < 0.05, \*\**P* < 0.01, \*\*\**P* < 0.001. Scale bars represent 1 cm (B), 2 mm (C) and 1 cm (E). Concentrations in μM are indicated in brackets.

We next aimed to characterize RN4 activity on lateral root development in these mutants. RN4 and RN4-1 reduced lateral root density in Col-0 and Ler (Fig. 5*B* and Fig S8*B*). Compared to Col-0, *axr5-1/iaa1* was resistant to RN4 at 5 μM, while *axr2-1/iaa7* was sensitive (Fig. 5*B*; Fig. S8*B*). Interestingly, *shy2-2/iaa3* was sensitive to RN4 at 5 μM, but resistant at 2 μM (Fig. 5*B* and Fig. S8*B*). Importantly, mutant sensitivity to RN4-1 was somewhat different to that observed with RN4. Our results suggest that AXR5/IAA1 might be degraded by both RN4-1 and RN4, while SHY2/IAA3 might be degraded specifically by RN4 and AXR2/IAA7 by RN4-1, to reduce lateral root density.

By using RN molecules together with their degradation products, we revealed potential contributions of specific AUX/IAAs to the complicated process of lateral root development. However, the sensitivities of the *aux/iaa* gain-of-function mutants to the RNs in terms of lateral root development did not exactly match the RN-induced AUX/IAA degradation/stabilization results found with our binding affinity assays. Lateral root development is a complicated process that requires the formation of a new meristem and emergence through several root layers, suggesting that the specific tissue context may affect RN activity and selectivity.

We therefore decided to switch our focus to apical hook development in etiolated seedlings, a rather simpler process than lateral rooting, but one also regulated by auxin (47). Apical hook development is characterized by differential growth between the two sides of the apical hypocotyl and comprises three phases named the formation, maintenance and opening phases (48–49). We first tested the effects of RN3 and RN4 on apical hook development in the wild type (Fig. S8*D*). While 2 μM RN3 did not affect apical hook development, RN4 completely abolished hook formation in a dose dependent manner (Fig. S8*D*; Fig. S8*E*).

We decided to exploit RN4 at 0.5 μM to understand whether selected AUX/IAAs play specific roles during apical hook development. We therefore tested the effects of 0.5 μM RN4 on hook development in the gain-of-function mutants *axr5-1/iaa1, axr2-1/iaa7* and *axr3-1/iaa17* for six days in the dark. All three mutants showed altered apical hook development compared to the wild type in control conditions (Fig. 5*C*, *E* and *G*). A detailed analysis of these results indicates that AXR5/IAA1 and AXR3/IAA17 need to be degraded for a proper apical hook to develop, while AXR2/IAA7 is likely stabilized during the formation phase and degraded during the maintenance phase. Similar to the wild type, *axr5-1*/*iaa1* showed sensitivity to RN4 during the formation phase, with no hook being present at 24 h; however, by 36 h the mutant had attained a slight hook curvature of 50 degrees, which then started opening directly (Fig. 5*D*). The mutant *axr2-1/iaa7* was resistant to RN4 in the formation phase (Fig. 5*F*) and *axr3-1/iaa17* was sensitive to RN4 (Fig. 5*H*). Taken together, these results indicate that all three AUX/IAAs tested here play a role during apical hook development. In particular, our results suggest that AXR2/IAA7 is stabilized during apical hook formation while AXR5/IAA1 stabilization occurs during the maintenance phase.

The effects of 0.5 μM RN4 on AUX/IAA mutants during the first 24 h of apical hook development (Fig. 5*D*, *F* and *H*) correlate strikingly with our *in vitro* pull-down assay results (Fig. 3*D*). AXR2/IAA7 proteins strongly interacted with TIR1 in the presence of RN4 (Fig. 3*D* and *G* and Fig. S6*B*), suggesting that a stabilized version of this AUX/IAA should confer resistance to the RN4 auxin agonist, which is indeed what we found with the *axr2-1/iaa7* gain-of-function mutant (Fig. 5*F*). In contrast, AXR5/IAA1 and AXR3/IAA17 did not interact with TIR1 when RN4 was present in the pull-down assay (Fig. 3*D*, *E* and *H* and Fig. S6*B*) and the corresponding gain-of-function mutants were sensitive to the effects of RN4 on hook development (Fig. 5*D* and *H*).

Overall, our study of the effects of the RN molecules on the AUX/IAA gain-of-function mutants distinguishes the involvement of specific AUX/IAAs in lateral root and apical hook development. Thus, we demonstrated the potential of such selective auxin agonists as the RNs in dissecting auxin perception controlling specific developmental processes *in vivo*.

### Mutation in the ATPase domain of *AtBRM* confers resistance to RN4

RN4 represents a useful tool to investigate the role of auxin during early stages of skotomorphogenesis. In order to identify new molecular players involved in apical hook development, we performed a forward genetic screen of sensitivity to RN4, using an EMS-mutagenized Col-0 population and selected those mutants that were able to form an apical hook in the presence of 0.5 μM RN4 in the dark, which we named *hookback (hkb)* mutants. We then further selected only those of the mutants that were sensitive to the effects of 75 nM 2,4-D on seedling phenotype in the light (Fig. S8*F*). Using this strategy, we could exclude known auxin resistant mutants that might appear in the screen. Several independent *hkb* lines, each carrying a single recessive mutation, were isolated from the screen and we focused on characterizing one of these, *hkb1*. In contrast to Col-0, *hkb1* had formed well-curved apical hooks in the presence of RN4 24 h after germination, while under mock-treated conditions there were no major differences between the two genotypes (Fig. 6*A*). Whole genome sequencing of *hkb1* revealed the presence of one non-synonymous EMS-like mutation (C to T nucleotide substitution) in the coding region of the *AT2G46020* gene that encodes for the SWItch/Sucrose Non-Fermentable (SWI/SNF) chromatin remodeling ATPase *BRAHMA (BRM)*. To confirm that the mutation in *BRM* is responsible for the resistance of *hkb1* against the negative effect of RN4 on apical hook formation, we carried out several analyses. First, we checked the phenotypes of available T-DNA mutants for *BRM*, including *brm-1, brm-2, brm-4* and *brm-5 (ectopic expression of seed storage proteins3, essp3)* (50–51). However, we focused our investigations on *brm-5* because both *hkb1* and *brm-5* contain a mutation in the ATPase domain (52) and 4-w-old plants of the two mutants showed similar phenotypes, including twisted leaves and less siliques than wild type (Fig. 6*B*). Importantly, *brm-5* showed similar resistance to the effect of 0.5 μM RN4 on apical hook formation to that shown by *hkb1* (Fig. 6*C* and *D*). These results strongly suggest that the mutation in the ATPase domain of *BRM* in *hkb1* is responsible for the resistance of this mutant to RN4. Next, we crossed *hkb-1* with *brm-5* and the F2 generation was analyzed. The *hkb1xbrm-5* mutant showed the same apical hook phenotype and similar RN4 resistance as the single *hkb1* and *brm-5* mutants (Fig. 6*C* and *D*), confirming that the mutation that confers resistance against RN4 in *hkb1* is in the *BRM* gene.

**Figure 6.**
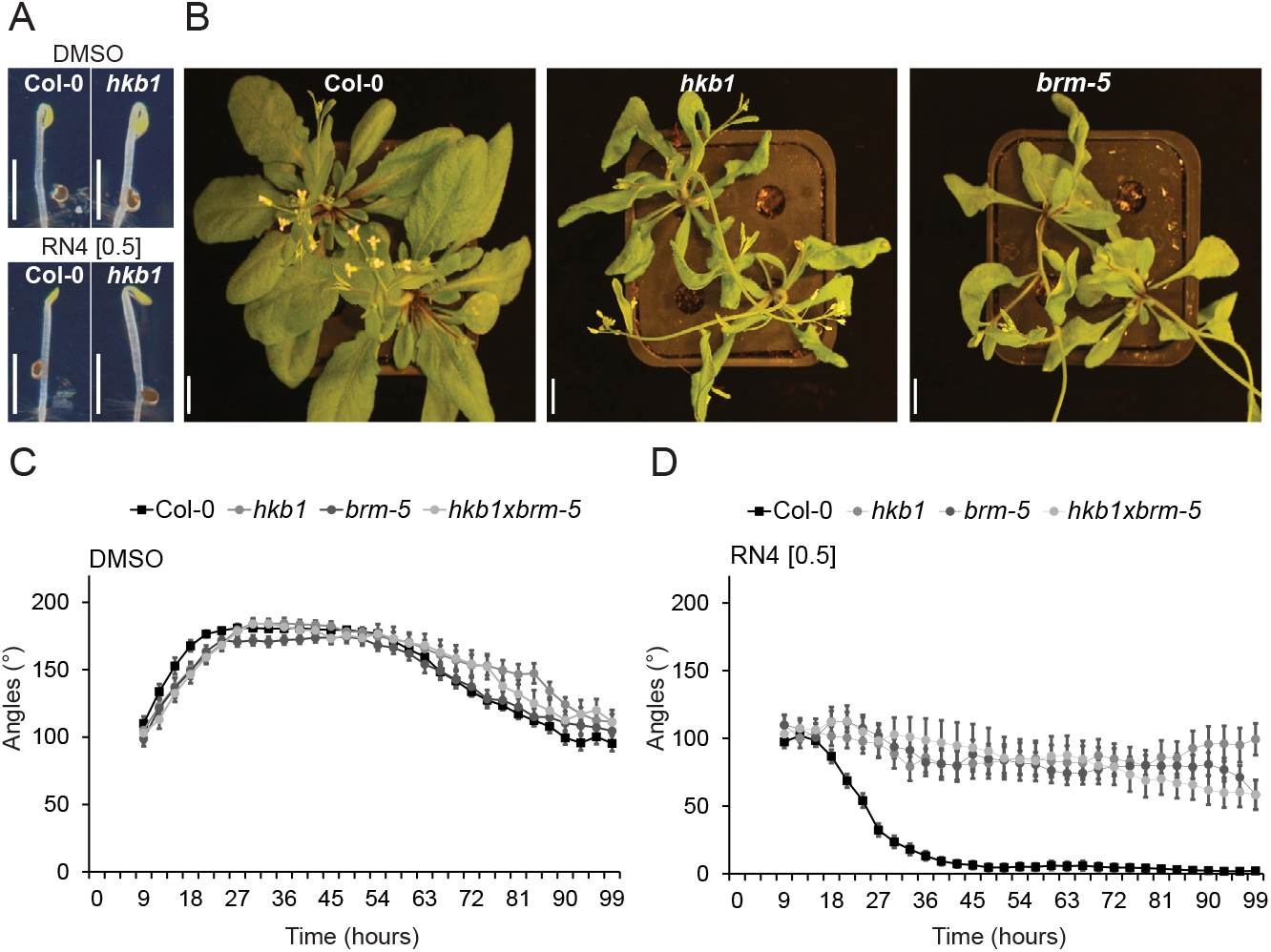
The *hkb1* mutant is resistant to the RN4 effect on apical hook development and carries a mutation on *BRAHMA (BRM)*. (A) Comparison of apical hook phenotype in Col-0 and *hkb1* seedlings 24 h after germination in the dark. The seedlings were grown on media supplemented with DMSO (upper panel) or RN4 (lower panel). While *hkb1* is resistant to the negative effect of RN4 on apical hook development, it appears similar to wild type in control conditions. (B) Four-w-old Col-0, *hkb1* and *brm-5* grown in long-day greenhouse conditions. (C-F) Apical hook angle in Col-0, *hkb1, brm-5* and *hkb1xbrm-5* (C-D) grown on DMSO (C) and 0.5 μM RN4 (D) supplemented media for 6 days in the dark. Measurement of apical hook angle was performed every three hours. Means ± SEM are shown, n > 18 seedlings across 2 independent replicates. Scale bars indicate 2 mm (A) and 1 cm (B). Concentrations in μM are indicated in brackets.

Our results suggest that BRM may function as a negative regulator of apical hook formation. Considering the resistance of both the *axr2/iaa7* gain-of-function mutant and *hkb1/brm-5* to the effect of RN4 on apical hook formation, we hypothesize that AXR2/IAA7 might negatively regulate BRM-induced gene transcription. We suggest that RN4 induces degradation of AXR2/IAA7, which may lead to BRM-mediated promotion of transcription of genes negatively regulating apical hook formation, potentially through chromatin remodeling.

Overall, our results show that selective auxin agonists can enable us to dissect the roles of specific AUX/IAAs in developmental processes, leading to the dissection of the molecular mechanisms of these processes.

## Discussion

Complicated auxin perception modules translate auxin signals into a multitude of developmental responses (53–54). Several studies have demonstrated that IAA displays different affinities for different SCF^TIR1/AFB^-AUX/IAA co-receptor complex combinations (6, 55) and specific auxin perception modules have even been shown to act sequentially during development (56). In this work, we discovered selective agonists of auxin and showed their potential to dissect these complex and redundant mechanisms that control selective aspects of plant development. We showed that specific auxin perception modules and their potential targets can be identified and characterized using the selective activities of auxin analogs such RN4. Remarkably, we even found variability of RN sensitivity between different accessions in both *Arabidopsis* and poplar, pointing to future challenges towards developing the most suitable auxin agonists for specific species and/or accessions.

Auxin behaves like molecular glue within the SCF^TIR1/AFB^-AUX/IAA co-receptor complex (53) by fitting into a space between the TIR1/AFB receptor and AUX/IAA co-receptor and extending the hydrophobic protein interaction surface. It has long been known that the auxin-binding pocket of SCF^TIR1/AFB^ is promiscuous, a feature which was heavily investigated during the early years of auxin research in the 1940s (57–58). During this time, several auxinic compounds were discovered including NAA, 2,4-D and picolinate auxins such as picloram (59), which are widely used today for basic research and agricultural applications. The 2,4-D and NAA modes of action are similar to that of IAA, as they also enhance the binding affinity between TIR1 and the AUX/IAAs. Their affinity to the co-receptor complex is lower than that of IAA, but both molecules are more stable metabolically, which explains their robust activity. Although the full details of the mode of action of these synthetic auxins are not yet known, these compounds have been instrumental in the discoveries of crucial auxin signaling components such as AXR1, AXR3/IAA17, AXR5/IAA1, AFB4 and AFB5 (60–64). Thus, synthetic compounds with auxin-like activities hold the potential to dissect the convoluted mechanisms of auxin signaling.

Here, we have shown the selective capacity of RN3 and RN4 to promote the interaction of TIR1 with specific AUX/IAA co-receptors, highlighting a strong potential for such auxin agonists in defining AUX/IAA involvement in specific transcriptional responses and developmental traits. This potential was strongly supported by our genetic approach, showing that different AUX/IAA gain-of-function mutants display defined sensitivities to RN3 and RN4 in terms of lateral root development. Multiple AUX/IAA-ARF modules act sequentially over time and space to orchestrate lateral root development (56, 65). Our data indicates that RN3 may promote development of lateral roots through SLR/IAA14 degradation and the stabilization of SHY2/IAA3, but the small number of *aux/iaa* gain-of-function mutants that we analyzed does not allow us to conclude whether degradation of additional AUX/IAAs is also required for this effect. On the other hand, the resistance of the *axr5-1/iaa1* mutant to high concentrations of RN4 reveals a novel role for the AXR5/IAA1 protein as a positive regulator of lateral root development.

Moreover, we could identify which of several AUX/IAA proteins are directly involved in apical hook development and reveal the implication of novel auxin-signaling components such as the SWI/SNF chromatin remodeling ATPase BRM. Remarkably, BRM has already been shown to be involved in auxin-dependent floral fate acquisition (66). In the inflorescence, when MONOPTEROS (MP)/ARF5 is free from AUX/IAA repression, it recruits BRM or its homolog SPLAYED (SYD) to remodel chromatin and thus promote gene transcription. Interestingly, in a yeast-three-hybrid assay, AXR3/IAA17 and BDL/IAA12 have been shown to prevent the association of MP to BRM (66). According to these results and our data showing the resistance of *axr2-1/iaa7* and *hkb1/brm-5* to RN4-mediated suppression of apical hook formation, we hypothesize that BRM, by associating with an unknown ARF transcription factor, might promote transcription of genes negatively regulating hook formation. We also hypothesize that AXR2/IAA7 might prevent the association of the ARF to BRM. Application of RN4 prompts the degradation of AXR2/IAA7, which may facilitate the association of the ARF to BRM, promoting transcription of downstream genes negatively regulating apical hook formation, potentially through chromatin remodeling. However, the hypothesis that stabilization of AXR2/IAA7 during apical hook formation blocks BRM activity raises the question of whether MP plays a role during hook development or whether BRM is recruited by other ARFs.

The different affinities of AUX/IAA proteins for IAA, RN3 and RN4 might lie in differences in residues within the DII domain. Our study thus brings us a step closer to a better quantitative understanding of the TIR1-AUX/IAA interaction system of auxin perception in a tissue-specific manner. Besides IAA, several other phytohormones including jasmonate-isoleucine, gibberellin, brassinosteroids and abscisic acid (ABA), also function by modulating the protein-protein interactions of their co-receptors (67). Isolation of novel molecules modulating the interactions of coreceptor complexes could therefore also be useful in uncovering the signaling components of these phytohormones.

Auxins have many uses in agriculture, horticulture, forestry and plant tissue culture (57). The selective auxin agonists described here may also find niche applications in these fields. RN activities in the low micromolar range and conservation of their specific developmental effects in land plants enforces this possibility. Moreover, the availability of models for ligand-bound co-receptors may allow rational design of a wider array of auxin agonists, using RN structures as a starting point. Indeed, this approach has already paved the way for developing agrochemicals interacting specifically with a subset of ABA receptors (69).

Overall, the isolation and characterization of chemical modulators of plant hormone signaling is an effective way to better understand the specificity of hormonal receptors. Because of the availability of genetic and genomic methods, most chemical biology approaches are performed in model species such as *Arabidopsis*. However, chemicals which induce well-characterized effects in *Arabidopsis* can be applied to non-model species to improve crop and tree value in agriculture and forestry, respectively. The complexity of the genomes of such non-model species may also be unraveled by the use of chemicals for which target proteins or pathways are known, giving a better understanding of evolutionary mechanisms.

## Materials and Methods

See *SI Materials and Methods* for detailed experimental procedures and additional materials and methods.

*Arabidopsis thaliana* seedlings were grown on ½ MS medium supplemented with 0.05% morpholinoethanesulfonic acid, 1% sucrose and 0.7% agar at pH 5.6. Stock solutions of all compounds used were dissolved in dimethyl sulfoxide (DMSO), which was also used in equal volume as a solvent control. Docking experiments were performed using SwissDock (69–70) with the ZINC ID of the RNs and 2P1Q crystal structure of TIR1 with the DII domain of AXR2/IAA7 (58). The best conformation was chosen according to the FullFitness (Kcal/mol). The corresponding binding energies for every conformation of each ligand were calculated using Hybrid-DFT-D3. The *in vitro* pull-down assays, with epitope-tagged TIR1 expressed with TnT-T7 coupled wheat germ extract (Promega), were performed as described previously (29, 71). For the luciferase assay, 7-day-old seedlings were incubated in Bright-Glo luciferase assay system (Promega) luciferine solution (LS) for 30 min before treatment with 50 μM compounds dissolved in LS. Light emission was recorded for 5 min at the indicated time point using a LAS-3000 (Fujifilm) and the natural log of the normalized relative light unit (RLU) was calculated as described previously (72). The degradation rate *k* (min^-1^) was used to compare the different treatments. The transcriptomic responses induced by the RNs were investigated by RNA-Seq, using *Arabidopsis thaliana* ecotype Col-0 cell suspension culture (73) treated with 50 μM RN3, RN4, or IAA for 30 min. Total RNA was extracted from filtered cells using the RNeasy Plant Mini Kit (QIAGEN) and sent to the SNP&SEQ Technology Platform in Uppsala University for sequencing. Genes were considered as being significantly differentially expressed if the adjusted p-values after FDR (False Discovery Rate) correction for multiple testing were lower than 0.05. For GUS assays, seedlings were fixed in 80% acetone, washed with 0.1 M phosphate buffer and transferred to 2 mM X-GlcA (Duchefa Biochemie) in GUS buffer (0.1 % triton X100; 10 mM EDTA; 0.5 mM potassium ferrocyanide; 0.5 mM potassium ferricyanide) in the dark at 37 °C before stopping the staining reaction with 70 % ethanol.

## Acknowledgments

We thank the many researchers who kindly provided us with published *Arabidopsis* lines and we acknowledge ABRC and NASC for distributing seeds. Sequencing was performed by the SNP&SEQ Technology Platform, Science for Life Laboratory at Uppsala University, a national infrastructure supported by the Swedish Research Council (VR-RFI) and the Knut and Alice Wallenberg Foundation. RNA-Seq data analysis was in part performed by BILS (Bioinformatics Infrastructure for Life Sciences). Amplification of *in vitro* poplar SwAsp lines was performed by the Poplar Transgenics Facility, Umeå Plant Science Centre. Whole genome resequencing was performed by Novogene and the results were analyzed with the support of Nicolas Delhomme and Iryna Shutava, Umeå Plant Science Centre Bioinformatics Platform. We would like to thank the Swedish Metabolomics Centre for access to instrumentation. We gratefully acknowledge O. Keech for critical reading of the manuscript. JC thanks J. Brown for technical assistance and K. Dreher and J. Gilkerson for helpful discussion and for transgenic line characterization. We thank M. Quareshy, V. Uzunova and R. Napier (University of Warwick) for assistance with insect cell expression of epitope-tagged TIR1. We are thankful to L. Bako who provided us with *Arabidopsis* cell culture, Pardeep Singh who shared powder of 2,4,5-T and RN4-1 with us and R. Bhalerao for sharing the system for time-lapse imaging of dark grown seedlings. This work was supported by Vetenskapsrådet VR 2013-4632 and VINNOVA (TV, ML, SD, P-AE, SR), Vetenskapsrådet VR 2016-00768 (QM), The Knut and Alice Wallenberg Foundation (AR), The Knut and Alice Wallenberg Foundation Shapesystem grant 2012-0050 (SD, SR, QM and KL), the Olle Engkvist Byggmästare Foundation (SaR), Kempestiftelserna (QM, DKB, P-AE), the Carl Tryggers Foundation (QM, MT), SweTree Technologies (SD), EMBO short term fellowship (TV), Seth M Kempe short term fellowship (TV), a travel grant from the Bröderna Edlunds Foundation (TV), National Science Foundation (MCB–0929100 to JC and ME), the Biotechnology and Biological Sciences Research Council (BB/L010623/1 to SK) and the Ministry of Education, Youth and Sports of the Czech Republic (the National Program for Sustainability I Nr. LO1204 to BP and ON). L.P. was funded by a postdoctoral scholarship and research grant 1507013N of the Research Foundation Flanders (FWO). Chemical Biology Consortium Sweden (CBCS) is primarily funded by the Swedish Research Council (P-AE).

## Footnotes

Author contributions: TV, LP, StR designed research; TV, SaR, NF, MK, QM, SMD, MT, BP, AR, MŁ performed research; NF, DKB, MK, BP, ON, AI, PAE, SRH, YZ, KL, JC, FA, SK, ME contributed new reagents or analytic tools; TV, SaR, NF, SMD, MT analyzed data; TV, SaR, SMD, StR wrote the paper.

The authors declare no conflict of interests.

